# External validation of machine learning models - registered models and adaptive sample splitting

**DOI:** 10.1101/2023.12.01.569626

**Authors:** Giuseppe Gallitto, Robert Englert, Balint Kincses, Raviteja Kotikalapudi, Jialin Li, Kevin Hoffschlag, Ulrike Bingel, Tamas Spisak

## Abstract

Multivariate predictive models play a crucial role in enhancing our understanding of complex biological systems and in developing innovative, replicable tools for translational medical research. However, the complexity of machine learning methods and extensive data pre-processing and feature engineering pipelines can lead to overfitting and poor generalizability. An unbiased evaluation of predictive models necessitates external validation, which involves testing the finalized model on independent data. Despite its importance, external validation is often neglected in practice due to the associated costs. Here we propose that, for maximal credibility, model discovery and external validation should be separated by the public disclosure (e.g. pre-registration) of feature processing steps and model weights. Furthermore, we introduce a novel approach to optimize the trade-off between efforts spent on training and external validation in such studies. We show on data involving more than 3000 participants from four different datasets that, for any “sample size budget”, the proposed adaptive splitting approach can successfully identify the optimal time to stop model discovery so that predictive performance is maximized without risking a low powered, and thus inconclusive, external validation. The proposed design and splitting approach (implemented in the Python package “AdaptiveSplit”) may contribute to addressing issues of replicability, effect size inflation and generalizability in predictive modeling studies.

## Introduction

Multivariate predictive models integrate information across multiple variables to construct predictions of a specific outcome and hold promise for delivering more accurate estimates than traditional univariate methods (Woo *et al*., 2017). For instance, in case of predicting individual behavioral and psychometric characteristics from brain data, such models can provide higher statistical power and better replicability, as compared to conventional mass-univariate analyses (Spisak *et al*., 2023). Predictive models can utilize a variety of algorithms, ranging from simple linear regression-based models to complex deep neural networks. With increasing model complexity, the model will be more prone to overfit its training dataset, resulting in biased, overly optimistic in-sample estimates of predictive performance and often decreased generalizability to data not seen during model fit (Hosseini *et al*., 2020)). Internal validation approaches, like cross-validation (cv) provide means for an unbiased evaluation of predictive performance during model discovery by repeatedly holding out parts of the discovery dataset for testing purposes (Efron & Tibshirani, 1994; Poldrack *et al*., 2020). However, internal validation approaches, in practice, still tend to yield overly optimistic performance estimates (Efron, 1983; Sui *et al*., 2020; Varoquaux & Cheplygina, 2022). There are several reasons for this kind of effect size inflation. First, predictive modelling approaches typically display a high level of “analytical flexibility” and pose a large number of possible methodological choices in terms of feature pre-processing and model architecture, which emerge as uncontrolled (e.g. not cross-validated) “hyperparameters” during model discovery. Seemingly ‘innocent’ adjustments of such parameters can also lead to overfitting, if it happens outside the cv loop. The second reason for inflated internally validated performance estimates is ‘leakage’ of information from the test dataset to the training dataset (Kapoor & Narayanan, 2023). Information leakage has many faces. It can be a consequence of, for instance, feature standardization in a non cv-compliant way or, in medical imaging, the co-registration of brain data to a study-specific template. Therefore, it is often very hard to notice, especially in complex workflows. Another reason for overly optimistic internal validation results may be that even the highest quality discovery datasets can only yield an imperfect representation of the real world. Therefore, predictive models might capitalize on associations that are specific to the dataset at hand and simply fail to generalize “out-of-the-distribution”, e.g. to different populations. Finally, some models might also be overly sensitive to unimportant characteristics of the training data, like subtle differences between batches of data acquisition or center-effects (Prosperi *et al*., 2020; Spisak, 2022).

The obvious solution for these problems is *external validation*; that is, to evaluate the model’s predictive performance on independent (‘external’) data that is guaranteed to be unseen during the whole model discovery procedure. There is a clear agreement in the community that external validation is critical for establishing machine learning model quality (Collins *et al*., 2014; Ho *et al*., 2020; Yu *et al*., 2022; Spisak *et al*., 2023; Poldrack *et al*., 2020). However, the amount of data to be used for model discovery and external validation can have crucial implications on the predictive power, replicability and validity of predictive models and is, therefore, subject of intense discussion (Riley *et al*., 2021; Marek *et al*., 2022; Spisak *et al*., 2023; Rosenberg & Finn, 2022; Thirion, 2023; Makowski *et al*., 2023; Supplementary Table 1). Finding the optimal sample sizes is especially challenging for biomedical research, where this trade-off needs to weigh-in ethical and economic considerations. As a consequence, to date only around 10% of predictive modeling studies include an external validation of the model (Yang *et al*., 2022). Those few studies performing true external validation often perform it on retrospective data (like Lee *et al*., 2021 or Kincses *et al*., 2023) or in separate, prospective studies (Spisak *et al*., 2020; Kincses *et al*., 2023). Both approaches can result in a suboptimal use of data and may slow down the dissemination process of new results.

In this manuscript we argue that maximal reliability and transparency during external validation can be achieved with prospective data acquisition preceded by “freezing” and publicly depositing (e.g. pre-registering) the whole feature processing workflow and all model weights. Furthermore, we present a novel adaptive design for predictive modeling studies with prospective data acquisition that optimizes the trade-off between efforts spent on training and external validation. We evaluate the proposed approach on data involving more than 3000 participants from four different datasets to illustrate that for any “sample size budget”, it can successfully identify the optimal time to stop model discovery, so that predictive performance is maximized without risking a low powered, and thus inconclusive, external validation.

## Background

### The anatomy of a prospective predictive modelling study

Let us consider the following scenario: a research group plans to involve a fixed number of participants in a study with the aim of constructing a predictive model, and at the same time, evaluate its external validity. How many participants should they allocate for model discovery, and how many for external validation, to get the highest performing model as well as conclusive validation results?

In most cases it is very hard to make an educated guess about the optimal split of the total sample size into discovery and external validation samples prior to data acquisition. A possible approach is to use simplistic rules-of-thumb. Splitting data with an 80-20% ratio (a.k.a Pareto-split, Lipovetsky, 2009) is probably the most common method, but a 90-10% or a 50-50% may also be plausible choices (Raykar & Saha, 2015). However, as illustrated on Figure 1, such prefixed sample sizes are likely sub-optimal in many cases and the optimal strategy is actually determined by the dependence of the model performance on training sample size, that is, the “learning curve”. For instance, in case of a significant but generally low model performance (Figure 1A: flat learning curve) the model does not benefit a lot from adding more data to the training set but, on the other hand, it may require a larger external validation set for conclusive evaluation, due to the lower predictive effect size. This is visualized by the “power curve” on Figure 1, which shows the statistical power of external validation with the remaining samples as a function of sample size used for model discovery. The optimal strategy will be different, however, if the learning curve shows a persistent increase, without a strong saturation effect, meaning that predictive performance can be significantly enhanced by training the model on larger sample size (Figure 1B). In this case, the stronger predictive performance that can be achieved with larger training sample size, at the same time, allows a smaller external validation sample to be still conclusive. Finally, in some situations, model performance may rapidly get strong and reach a plateau at a relatively low sample size (Figure 1C). In such cases, the optimal strategy might be to stop early with the discovery phase and allocate resources for a more powerful external validation.

**Figure 1:**
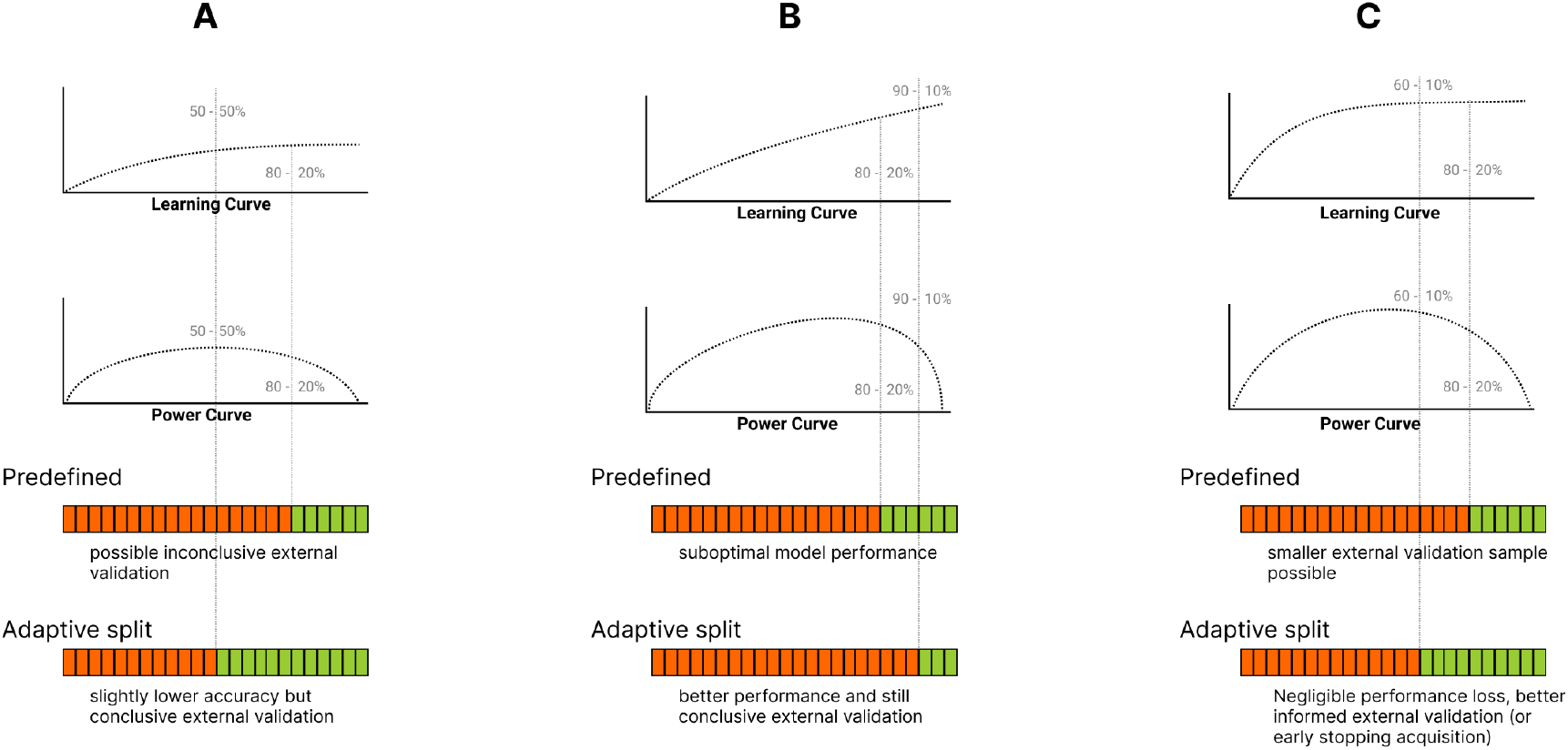
Examples of different optimal discovery and external validation sample sizes compared to a predefined 80-20% Pareto-split. **(A)** If the planned sample size and the model performance is low, the predefined external validation sample size might provide low statistical power to detect a significant model performance. **(B)** External validation of highly accurate models is well-powered; increasing the training sample size (against the external validation sample size) might result in a better performing final model. **(C)** Continuing training on the plateau of the learning curve will result in a negligible or biologically not relevant model performance improvement. In this case, a larger external validation sample (for more robust external performance estimates) or ‘early stopping’ of the data acquisition process might be desirable.

### Transparent reporting of external validation: registered models

A key criterion for external validation is the independence of the external data from the data used during model discovery (Steyerberg & Harrell, 2016; Collins *et al*., 2014; Spisak *et al*., 2023). Regardless of the splitting strategy, an externally validated predictive modelling study must provide strong guarantees for this independence criterion. Pre-registration, i.e. the public disclosure of study plans before the start of the study, is an increasingly popular way of enhancing transparency and replicability in biomedical research (Nosek *et al*., 2019; Spisak *et al*., 2023) (Figure 2A), which could also be used to ensure the independence of the external validation sample.

**Figure 2:**
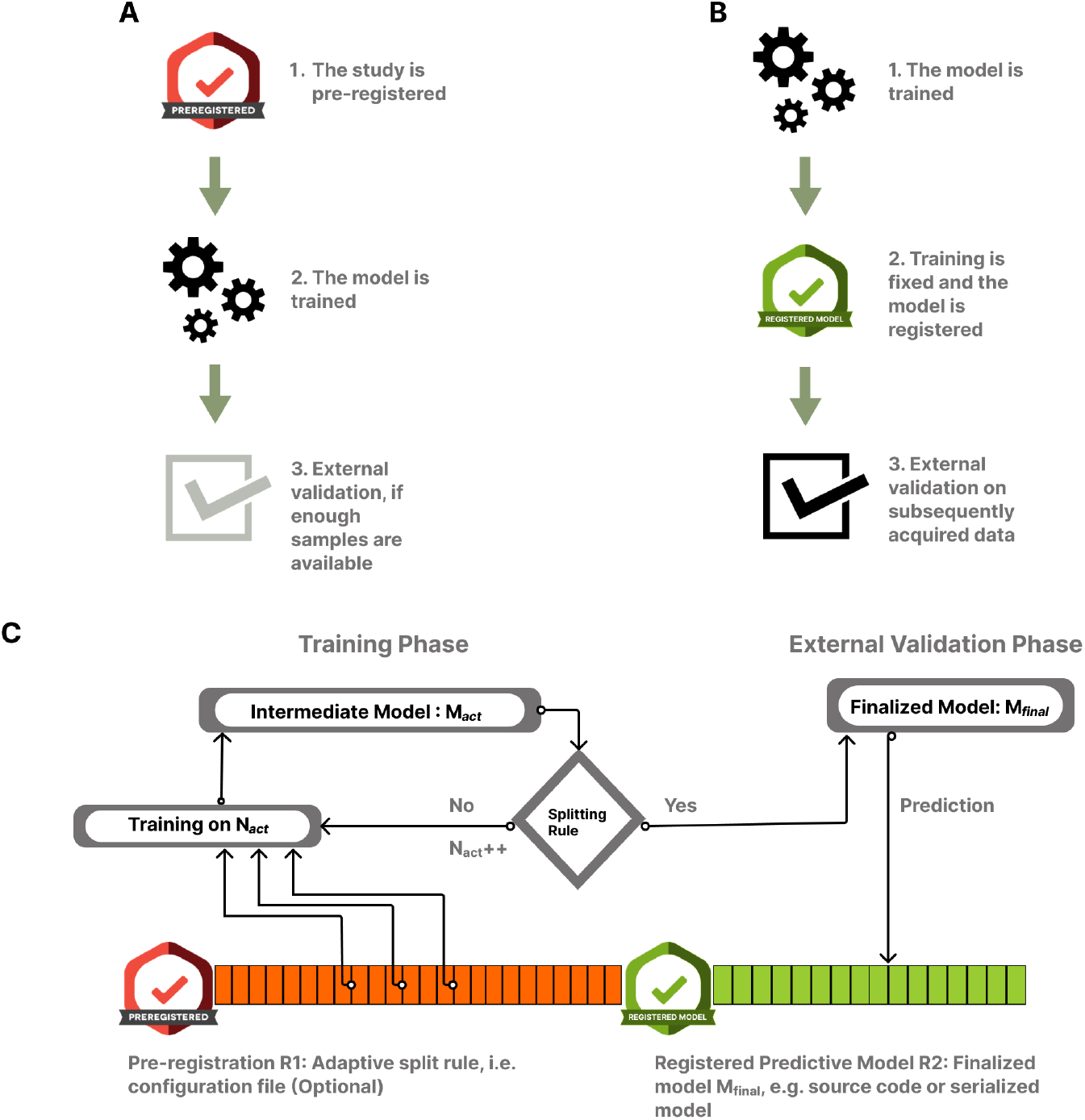
The registered model design and the proposed adaptive sample splitting procedure for prospective predictive modeling studies. **(A)** Predictive modelling combined with conventional pre-registration. In this case the pre-registration precedes data acquisition and requires fixing as many details of the analysis as possible. Given the potentially large number of coefficients to be optimized and the importance of hyperparameter optimization, conventional pre-registration exhibits a limited compatibility with predictive modelling studies. **(B)** Here we propose that in case of predictive modelling studies, public registration should only happen after the model is trained and finalized. The registration step in this case includes publicly depositing the finalized model, with all its parameters as well as all feature pre-processing steps. External validation is performed with the resulting *registered model*. This practice ensures a transparent, clear separation of model discovery and external validation. **(C)** The “registered model” design allows a flexible, adaptive splitting of the “sample size budget” into discovery and external validation phases. The proposed adaptive sample splitting procedure starts with fixing (and potentially pre-registering) a stopping rule (R1). During the training phase, one or more candidate models are trained and the splitting rule is repeatedly evaluated as the data acquisition proceeds. When the splitting rule “activates”, the model gets finalized (e.g. by being fit on the whole training sample) and publicly deposited/registered (R2). Finally, data acquisition continues and the prospective external validation is performed on the newly acquired data.

However, as the concept of pre-registration was originally developed for confirmatory research, it does not fit well with the exploratory nature of the model discovery phase in typical predictive modelling endeavors. Specifically, while pre-registration necessitates that as many parameters of the analysis as possible are fixed before data acquisition, predictive modelling studies often involve a large number of hyperparameters (e.g. model architecture, feature pre-processing steps, regularization parameters, etc.) that are not known in advance and need to be optimized during the model discovery phase. This is especially true for complex machine learning models, like deep neural networks, where the number of free parameters can easily reach tens of thousands or even more. In such cases, the pre-registration of the discovery phase would require a large number of assumptions or simplifications, which would make the process ineffective and less transparent.

Therefore, we propose to perform the pre-registration after the model discovery phase, but before the external validation (Figure 2B). In this case, more freedom is granted for the discovery phase, while the external validation remains equally conclusive, as long as the pre-registration of the external validation includes all details of the *finalized* model (including the feature pre-processing workflow). This can easily be done by attaching the data and the reproducible analysis code used during the discovery phase or, alternatively, a serialized version of the fitted model (i.e. a file that contains all model weight, see e.g. Spisak *et al*., 2020 and Kincses *et al*., 2023). We refer to such models as **registered models**.

### The adaptive splitting design

Even with registered models, the amount of data to be used for model discovery and external validation can have crucial implications on the predictive power, replicability and validity of predictive models. Here, we introduce a novel design for prospective predictive modeling studies that leverages the flexibility of model discovery granted by the registered model design. Our approach aims to adaptively determine an optimal splitting strategy during data acquisition. This strategy balances the model performance and the statistical power of the external validation (Figure 2C). The proposed design involves continuous model fitting and hyperparameter tuning throughout the discovery phase, for example, after every 10 new participants, and evaluating a ‘stopping rule’ to determine if the desired compromise between model performance and statistical power of the external validation has been achieved. This marks the end of the discovery phase and the start of the external validation phase, as well as the point at which the model must be publicly and transparently deposited or preregistered. Importantly, the preregistration should precede the continuation of data acquisition, i.e., the start of the external validation phase. In the present work, we propose and evaluate a concrete, customizable implementation for the splitting rule.

## Methods and Implementation

### Components of the stopping rule

The stopping rule of the proposed adaptive splitting design can be formalized as function *S*:

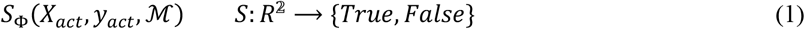

where Φ denotes customizable parameters of the rule (detailed in the next paragraph), *X*_*act*_ ∈ *R*^𝟚^ is the data (a matrix consisting of *n*_*act*_ > 0 observations and a fixed number of features *p*) and *y*_*act*_ ∈ *R* is the prediction target, as acquired so far and *ℳ* is the machine learning model to be trained. The discovery phase ends if and only if the stopping rule returns *True*.

#### Hard sample size thresholds

Our stopping rule is designed so that it can force a minimum size for both the discovery and the external validation samples, *t*_*min*_ and *v*_*min*_, both being free parameters of the stopping rule.

Specifically:

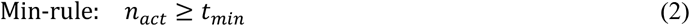

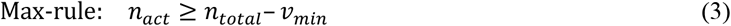

where *n*_*act*_ and *n*_*total*_ are the actual sample size (e.g. participants measured so far) and the total sample size (i.e. the “sample size budget”), respectively, so that *n*_*total*_ ≥ *n*_*act*_ > 0. Setting *t*_*min*_ and *v*_*min*_ may be useful to prevent early stopping at the beginning of the training procedure, where predictive performance and validation power estimates are not yet reliable due to the small *n*_*act*_ or to ensure that a minimal validation sample size, even if stopping criteria are never met. If *t*_*min*_ and *v*_*min*_ are set so that *t*_*min*_ + *v*_*min*_ = *n*_*total*_ then our approach falls back to training a registered model with predefined training and validation sample sizes.

#### Forecasting Predictive Performance via Learning Curve Analysis

Taking internally validated performance estimates of the candidate model as a function of training sample size, also known as learning curve analysis, is a widely used approach to gain deeper insights into model training dynamics (see examples on Figure 1). In the proposed stopping rule, we will rely on learning curve analysis to provide estimates of the current predictive performance and the expected gain when adding new data to the discovery sample.

Performance estimates can be unreliable or noisy in many cases, for instance with low sample sizes or when using leave-one-out cross-validation (Varoquaux, 2018). To obtain stable and reliable learning curves, we propose to calculate multiple cross-validated performance estimates from sub-samples sampled without replacement from the actual data set. The proposed procedure is detailed in Algorithm 1.

##### Algorithm 1

**(Bootstrapped Learning Curve Analysis)**

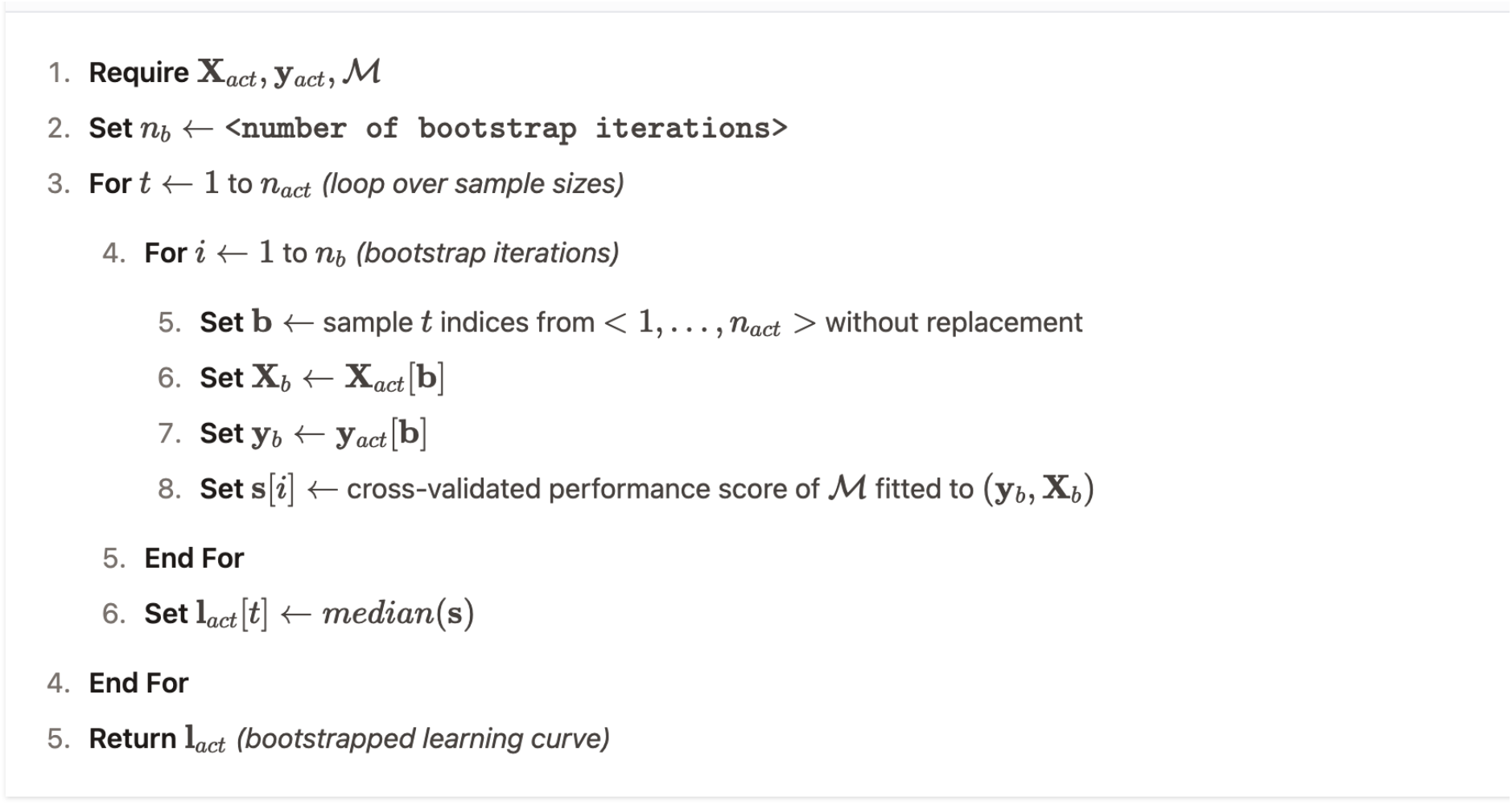

The learning curve analysis allows the discovery phase to be stopped if the expected gain in predictive performance is lower than a predefined relevance threshold and can be used for instance for stopping model training earlier in well-powered experiments and retain more data for the external validation phase. Specifically, the stopping rule *S* will return *True* if the *Min-rule* (Eq. 2) is *True* or the following is true:

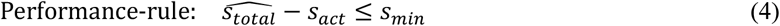

where *s*_*act*_ is the actual bootstrapped predictive performance score (i.e. the last element of *l*_*act*_, as returned by Algorithm 1, 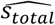 is a estimate of the (unknown) predictive performance *s*_*total*_ (i.e. the predictive performance of the model trained on the whole sample size) and ϵ_*s*_ is the smallest predictive effect of interest. Note that setting ϵ_*s*_ = −∞ deactivates the *Performance-rule* (Eq. 4).

While *s*_*total*_ is typically unknown at the time of evaluating the stopping rule *S*, there are various approaches of obtaining an estimate 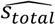. In the base implementation of AdaptiveSplit, we stick to a simple method: we extrapolate the learning curve *l*_*act*_ based on its tangent line at *n*_*act*_, i.e. assuming that the latest growth rate will remain constant for the remaining samples. While in most scenarios this is an overly optimistic estimate, it still provides a useful upper bound for the maximally achievable predictive performance with the given sample size and can successfully detect if the learning curve has already reached a flat plateau (like on Figure 1C).

#### Statistical power of the external validation sample

Even if the learning curve did not reach a plateau, we still need to make sure that we stop the training phase early enough to save a sufficient amount of data for a successful external validation from our sample size budget. Given the actual predictive performance estimate *s*_*act*_ and the size of the remaining, to-be-acquired sample *s*_*total*_ − *s*_*act*_, we can estimate the probability that the external validation correctly rejects the null hypothesis (i.e. zero predictive performance). This type of analysis, known as power calculation, allows us to determine the optimal stopping point that guarantees the desired statistical power during the external validation. Specifically, the stopping rule *S* will return *True* if the *Performance-rule* (Eq. 4) is *False* and the following is true:

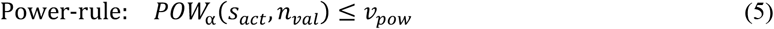

where *POW*_α_ (*s, n*) *no* is the power of a validation sample of size *n* to detect an effect size of *s* and *n*_*val*_ = *n*_*total*_ − *n*_*act*_ is the size of the validation sample if stopping, i.e. the number of remaining (not yet measured) participants in the experiment. Given that machine learning model predictions are often non-normally distributed (Spisak, 2022), our implementation is based on a bootstrapped power analysis for permutation tests, as shown in Algorithm 2. Our implementation is, however, simple to extend with other parametric or non-parametric power calculation techniques.

##### Algorithm 2

**(Calculating of the power-rule)**

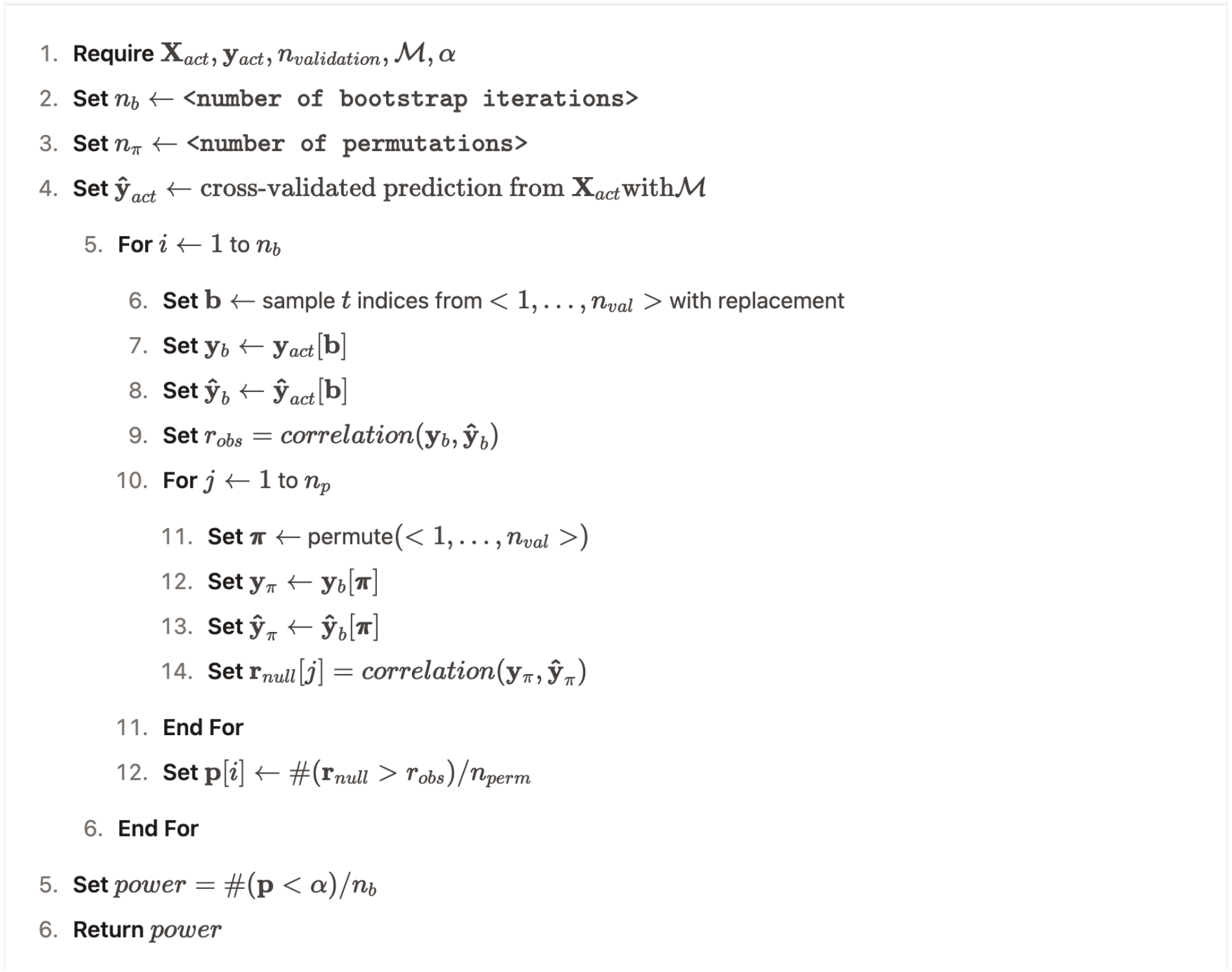

Note that depending on the aim of external validation, the *Power-rule* can be swapped to, or extended with, other conditions. For instance, if we are interested in accurately estimating the predictive effect size, we could condition the stopping rule on the width of the confidence interval for the prediction performance.

Calculating the validation power (Algorithm 2) for all available sample sizes (*n* = 1 … *n*_*act*_) defines the so-called “validation power curve” (see Figure 1 and Supplementary Figures 2, 4 and 6), that represents the expected ratio of true positive statistical tests on increasing sample size calculated on the external validation set. Various extrapolations of the power curve can predict the expected stopping point during the course of the experiment.

### Stopping Rule

Our proposed stopping rule integrates the Min*-*rule, the Max*-*rule, the Performance*-*rule and the Power*-*rule in the following way:

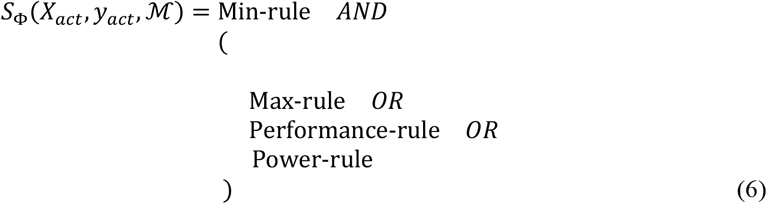

where Φ =< *t*_*min*_, *v*_*min*_, *s*_*min*_, *v*_*pow*_, α > are parameters of the stopping rule: minimum training sample size, minimum validation sample size, minimum effect of interest and target power for the external validation and the significance threshold, respectively.

We have implemented the proposed stopping rule in the Python package “*adaptivesplit*^1^. The package can be used together with a wide variety of machine learning tools and provides an easy-to-use interface to work with scikit-learn (Pedregosa *et al*., 2012) models.

### Empirical evaluation

We evaluate the proposed stopping rule, as implemented in the package *adaptivesplit*, in four publicly available datasets; the Autism Brain Imaging Data Exchange (ABIDE; Di Martino *et al*., 2013), the Human Connectome Project (HCP; Van Essen *et al*., 2013), the Information eXtraction from Images (IXI)^2^ and the Breast Cancer Wisconsin (BCW; Street *et al*., 1993) datasets (Fig. 3).

**Figure 3:**
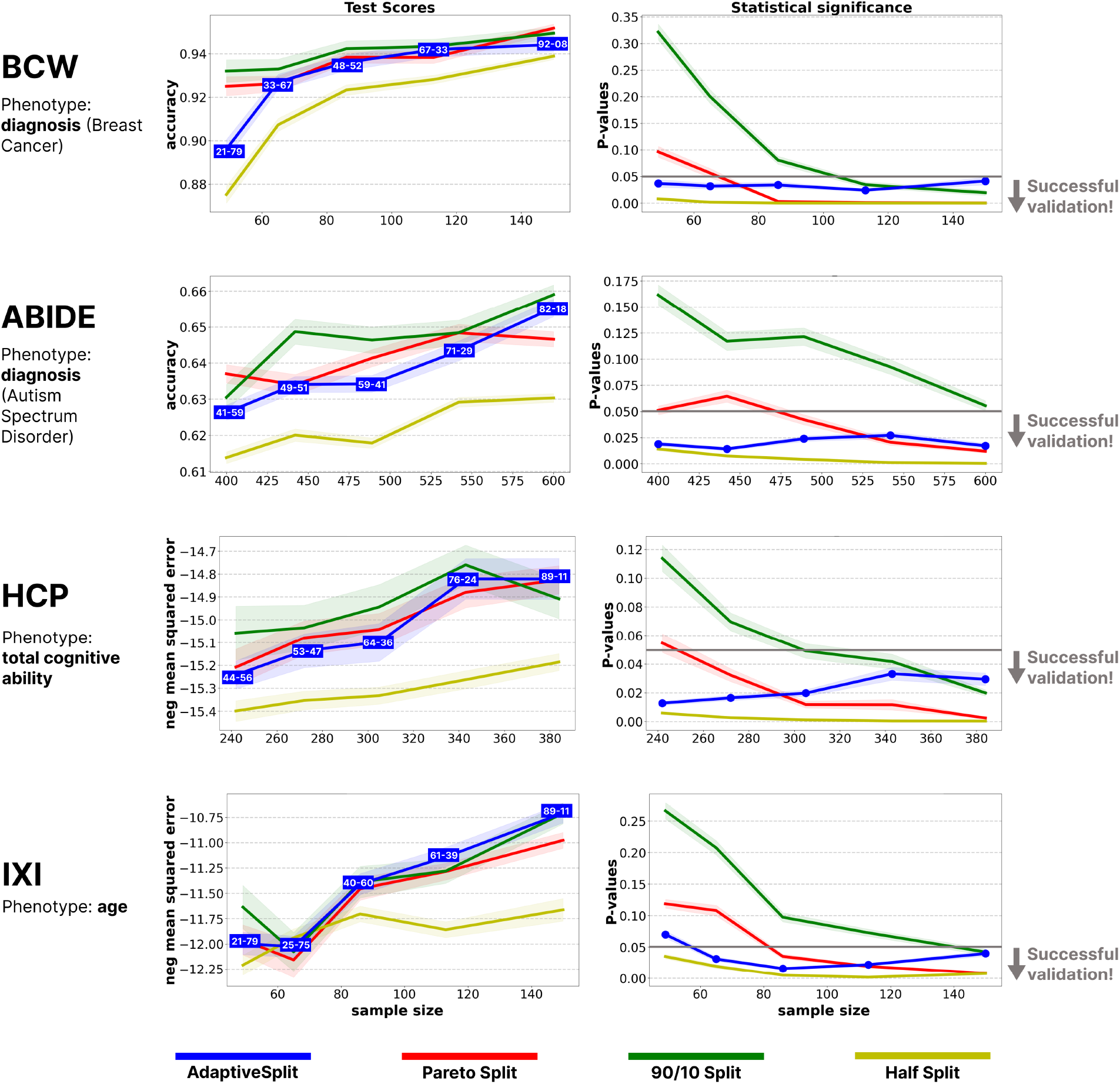
The proposed adaptive splitting approach provides a good compromise between predictive performance and statistical power of the external validation. The left and right column shows the comparison of splitting methods on external validation performance and p-values, respectively, at various *n*_*total*_. Confidence intervals are based on 100 repetitions of the analyses. The adaptive splitting approach (blue) provides a good compromise between predictive performance and statistical power of the external validation. The Pareto split (orange) provides similar external validation performances to adaptive splitting; however it often fails to provide conclusive results due to an insufficient sample size during external validation, especially in case of a limited sample size budget. The 90-10% split (green) provides only slightly higher performances than the Pareto and the Adaptive splitting techniques, but it very often gives inconclusive results (*p* ≥ 0.05) in the external validation sample. Half-split (red) tends to provide worse predictive performance due to the too small training sample.

#### ABIDE

We obtained preprocessed data from Autism Brain Imaging Data Exchange (ABIDE) dataset (Di Martino *et al*., 2013) involving 866 participants (Autism Spectrum Disorder: 402, neurotypical control: 464). Pre-processed regional time-series data were obtained as shared^3^ by Dadi *et al*., 2019, which were based on pre-processed image data provided by the Pre-processed Connectome Project (Cameron *et al*., 2013). Tangent correlation across the time series of the n=122 regions of the BASC brain parcellation (Multi-level bootstrap analysis of stable clusters; Bellec *et al*., 2010) was computed with nilearn^4^. The resulting functional connectivity estimates were considered features for a predictive model of autism diagnosis.

#### HCP

The Human Connectome Project dataset contains imaging and behavioral data of approximately 1,200 healthy subjects (Van Essen *et al*., 2013). Pre-processed resting state functional magnetic resonance imaging (fMRI) connectivity data (partial correlation matrices; Glasser *et al*., 2013 as published with the HCP1200 release (N=999 participants with functional connectivity data) were used to build models that predict individual fluid intelligence scores (Gf), measured with Penn Progressive Matrices (Duncan *et al*., 2000).

#### IXI

The IXI dataset is published by the Neuroimage Analysis Center, from Imperial College London, in the United Kingdom, and it is part of the project Brain Development. It consists of approximately 600 structural MRI images from a diverse population of healthy individuals, including both males and females across a wide age range. The dataset contains high-resolution brain images from three different MRI scanners (Philips Intera 3T, Philips Gyroscan Intera 1.5T and GE 1.5T) and associated demographic information, making it suitable for studying age-related changes in brain structure and function. We used gray matter probability maps generated from T1– weighted MR images with Freesurfer (Fischl, 2012) as features for a predictive model of age.

#### BCW

The Breast Cancer Wisconsin (BCW, Street *et al*., 1993) dataset contains diagnostic features computed from a digitized image of a fine needle aspirate (FNA) of a breast mass. The dataset includes 30 different features such as the mean radius, mean texture, mean perimeter, mean area, mean smoothness, mean compactness, mean concavity, etc. The target variable for predictive modelling in this dataset is the diagnosis (M = malignant, B = benign).

The chosen datasets include both classification and regression tasks, and span a wide range in terms of number of participants, number of predictive features, achievable predictive effect size and data homogeneity (see Supplementary Figures 1-6). Our analyses aimed to contrast the proposed adaptive splitting method with the application of fixed training and validation sample sizes, specifically using 50, 60 or 90% of the total sample size for training and the rest for external validation. We simulated various “sample size budgets” (total sample sizes, *n*_*total*_) with random sampling without replacement. For a given total sample size, we simulated the prospective data acquisition procedure by incrementing *n*_*act*_; starting with 10% of the total sample size and going up with increments of five. In each step, the stopping rule was evaluated with “AdaptiveSplit”, fitting a Ridge model (for regression tasks) or a L2-regularized logistic regression (for classification tasks). Model fit always consisted of a cross-validated fine-tuning of the regularization parameter, resulting in a nested cv estimate of prediction performance and validation power. Robust estimates (and confidence intervals) were obtained with bootstrapping, as described in Algorithm 1 and Algorithm 2. This procedure was iterated until the stopping rule returned True. The corresponding sample size was then considered the final training sample. With all four splitting approaches (adaptive, Pareto, Half-split, 90-10% split), we trained the previously described Ridge or regularized logistic regression model on the training sample and obtained predictions for the sample left out for external validation. This whole procedure was repeated 100 times for each simulated sample size budget in each dataset, to estimate the confidence intervals for the models performance in the external validation and its statistical significance. In all analyses, the adaptive splitting procedure is performed with a target power of *v*_*pow*_ = 0.8, an *alpha* = 0.05, *t*_*tmin*_ = *n*_*total*_/3, *v*_*min*_ = 12, *s*_*min*_ = −∞. P-values were calculated using a permutation test with 5000 permutations.

## Results

The results of our empirical analyses of four large, openly available datasets confirmed that the proposed adaptive splitting approach can successfully identify the optimal time to stop acquiring data for training and maintain a good compromise between maximizing both predictive performance and external validation power with any sample size budget.

In all four samples, the applied models yielded a statistically significant predictive performance at much lower sample sizes than the total size of the dataset, i.e. all datasets were well powered for the analysis. Trained on the full sample size with cross-validation, the models displayed the following performances: functional brain connectivity from the HCP dataset explained 13% of the variance in cognitive abilities; structural MRI data (gray matter probability maps) in the IXI dataset explained 48% in age; classification accuracy was 65.5% for autism diagnosis (functional brain connectivity) in the ABIDE dataset and 92% for breast cancer diagnosis in the BCW dataset.

The datasets varied not only in the achievable predictive performance but also in the shape of the learning curve, with different sample sizes and thus, they provided a good opportunity to evaluate the performance of our stopping rule in various circumstances (Supplementary Figures 1-6).

We found that adaptively splitting the data provided external validation performances that were comparable to the commonly used Pareto split (80-20%) in most cases (Figure 3, left column). As expected half-split tended to provide worse predictive performance due to the smaller training sample. In contrast, 90-10% tended to display only slightly higher performances than the Pareto and the Adaptive splitting techniques, in most cases. This small achievement came with a big cost in terms of the statistical power in the external validation sample, where the 90-10% split very often gave inconclusive results (*p* ≥ 0.05) (Figure 3, right column), especially with low sample size budgets. Although to a lesser degree, Pareto split also frequently failed to yield a conclusive external validation with small total sample sizes. Adaptive splitting (as well as half-split) provided sufficient statistical power for the external validation in most cases. This was achieved by applying different strategies in different scenarios. In case of low total sample sizes, it retained a larger proportion of the sample for the external validation phase in order to achieve sufficient power, up to using 79% of the data for external validation. On the other hand, if the total sample size budget allowed it, adaptive splitting let the predictive model benefit from larger training samples, retaining as low as 8% of the data for external validation is such cases.

Focusing only on cases with a successful, conclusive external validation, the proposed adaptive splitting strategy always provided equally good or better predictive performance than the fixed splitting strategies (as shown by the 95% confidence intervals on Figure 3).

## Discussion

Here we have proposed “registered models”, a novel design for prospective predictive modeling studies that allows flexible model discovery and trustworthy prospective external validation by fixing and publicly depositing the model after the discovery phase. Furthermore, capitalizing on the flexibility during model discovery with the registered model design, we have proposed a stopping rule for adaptively splitting the sample size budget into discovery and external validation phases. These approaches together provide a robust and flexible framework for predictive modeling studies and address several common issues in the field, including overfitting, effect size inflation as well as the lack of reliability and reproducibility.

Registered models provide a clear and transparent separation between the discovery and external validation phases, which is essential for ensuring the independence of the external validation data. Thereby, they provide a straightforward solution to several of the widely discussed issues and pitfalls of predictive model development (Efron, 1983; Sui *et al*., 2020; Varoquaux & Cheplygina, 2022; Marek *et al*., 2022; Spisak *et al*., 2023). With registered models, external validation estimates are guaranteed to be free of information leakage (Kapoor & Narayanan, 2023) and provide an unbiased estimate of the model’s predictive performance.

With registered models, the question of how the total sample size budget should be distributed between the discovery and external validation phase remains of central importance for the optimal use of available resources (scanning time, budget, limitations in participant recruitment) (Archer *et al*., 2020; Riley *et al*., 2021; Marek *et al*., 2022; Spisak *et al*., 2023; Rosenberg & Finn, 2022; Thirion, 2023; Makowski *et al*., 2023; Supplementary Table 1). Optimal sample sizes are often challenging to determine prior to the study. The proposed adaptive splitting procedure promises to provide a solution in such cases by allowing the sample size to be adjusted during the data acquisition process, based on the observed performance of the model trained on the already available data. We performed a thorough evaluation of the proposed adaptive splitting procedure on data from more than 3000 participants from four publicly available datasets. We found that the proposed adaptive splitting approach can successfully identify the optimal time to stop acquiring data for training and maintain a good compromise between maximizing both predictive performance and external validation power with any “sample size budget”. When contrasting splitting approaches based on fixed validation size with the proposed adaptive splitting technique, using the latter was always the preferable strategy to maximize power and statistical significance during external validation. The benefit of adaptively splitting the data acquisition for training and validation provides the largest benefit in lower sample size regimes. In case of larger total sample size budgets, the fixed Pareto split (20-80%) provided also good results, giving similar external validation performances to adaptive splitting, without having to repeatedly re-train the model during data acquisition. Thus, for moderate to large sample sizes and well powered models, the Pareto split might be a good alternative to the adaptive splitting approach, especially if the computational resources for re-training the model are limited.

Of note, the presented implementation of adaptive data splitting aims to maximize the training sample (and minimize the external validation sample) in order to achieve the highest possible performance together with a conclusive (statistically significant) external validation. However, the resulting external performance estimates will still be subject of sampling variance. If the aim is to provide more reliable estimates of the predictive effect size in the external validation, the power-rule in the proposed approach can be modified so that it stops the discovery phase when a desired confidence interval width for the external effect size estimate is reached.

The proposed adaptive splitting design can advance the development of predictive models in several ways. Firstly, it provides a simple way to perform both model discovery and initial external validation in a single study. Furthermore, it promotes the public deposition (registration) of models at an early stage of the study, enhancing transparency, reliability and replicability. Finally, it provides a flexible approach to data splitting, which can be adjusted according to the specific needs of the study.

In conclusion, registered models provide a simple approach to guarantee the independence of model discovery and external validation and for the development and initial evaluation of registered models with unknown power, the introduced adaptive splitting procedure provides a robust and flexible approach to determine the optimal ratio of data to be used for model discovery and external validation. Together, registered models and the adaptive splitting procedure, address several common issues in the field, including overfitting, cross-validation failure, and boost the reliability and reproducibility.

## Acknowledgements

The work is funded by the Deutsche Forschungsgemeinschaft (DFG, German Research Foundation) - Project-ID 422744262 - TRR 289 (Gefördert durch die Deutsche Forschungsgemeinschaft (DFG) – Projektnummer 422744262 – TRR 289).

## Footnotes

1 https://github.com/pni-lab/adaptivesplit

2 https://brain-development.org/ixi-dataset/

3 https://osf.io/hc4md

4 http://nilearn.github.io/

## Notes

https://pni-lab.github.io/adaptivesplit/supplementary

### Competing Interest Statement

The authors have declared no competing interest.

### Summary of Updates

1. An author name missing from the old version of the manuscript has been added 2. Fixed typos in the captions for Fig.1 and Fig.2, in the supplemental material 3. hyphens added in the main text for consistency purposes (e.g. preprocessing -< pre-processing) 4. Acknowledgements section added 5. New references added 6. Abstract modified for better text flow 7. Small changes to main text for better flow 8. Size of figure 2 and figure 3 revised.

