## Supplementary Material for "External validation of machine learning models - registered models and adaptive sample splitting"

#### Supplementary Figures

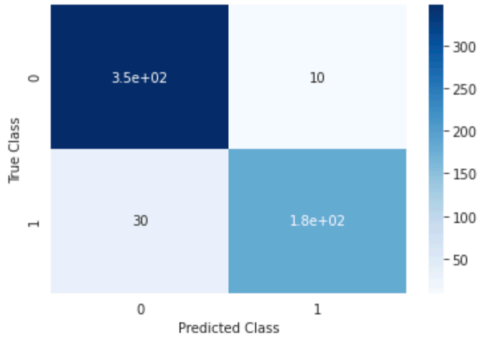


**Figure 1:** Predictive performance (confusion matrix) of the model trained on the BCW dataset to predict diagnosis. The model was trained on the whole dataset with nested cross-validation.

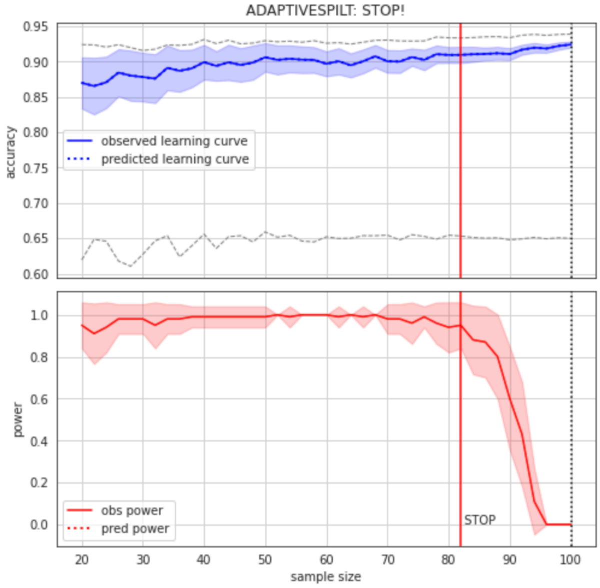


**Figure 2:** Learning curve (top) and power curve (bottom) of the model trained on the BCW dataset to predict diagnosis. The maximum sample size (i.e. the whole dataset) was considered as the "sample size budget". X-axis: $n_{act}$; y-axis (learning curve): Accuracy as a measure of predictive performance; y-axis (power curve): statistical power of the remaining sample to confirm the model's validity.

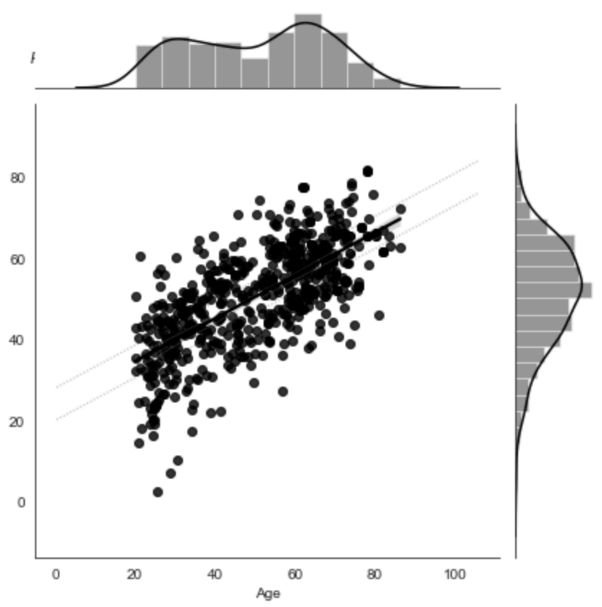


**Figure 3:** Predictive performance of the model trained on gray matter probability images from the IXI dataset to predict age. The model was trained on the whole dataset with nested cross-validation. X-axis: true age, y-axis: predicted age.

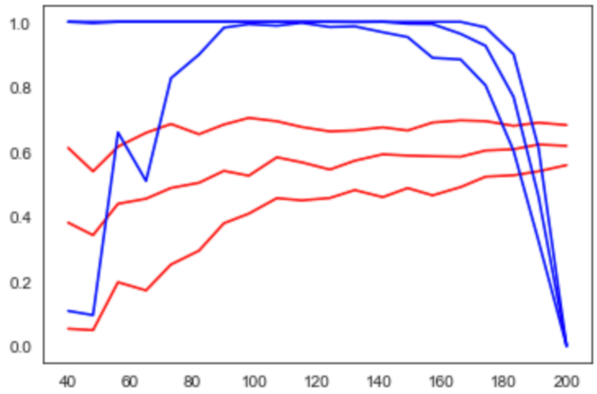


**Figure 4:** Learning curve (red) and power curve (blue) of the model trained on gray matter probability images from the IXI dataset to predict age. The maximum sample size (i.e. the whole dataset) was considered as the "sample size budget". X-axis: $n_{act}$; y-axis (learning curve): Pearson's correlation as a measure of predictive performance; y-axis (power curve): statistical power of the remaining sample to confirm the model's validity.

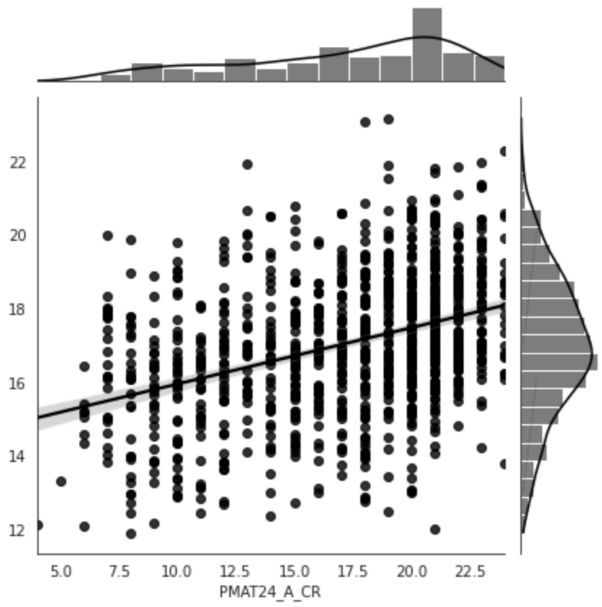


**Figure 5:** Predictive performance of the model trained on resting state functional connectivity data from the HCP dataset to predict fluid intelligence (PMAT24_A_CR). The model was trained on the whole dataset with nested cross-validation. X-axis: true age, y-axis: predicted age.

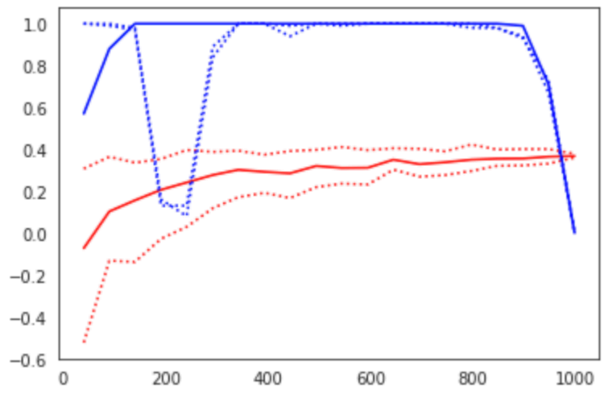


**Figure 6:** Learning curve (red) and power curve (blue) of the model trained on resting state functional connectivity data from the HCP dataset to predict fluid intelligence (PMAT24_A_CR). The maximum sample size (i.e. the whole dataset) was considered as the "sample size budget". X-axis: $n_{act}$; y-axis (learning curve): Pearson's correlation as a measure of predictive performance; y-axis (power curve): statistical power of the remaining sample to confirm the model's validity.

### Supplementary Tables

#### Supplementary Table 1

Manuscripts, commentaries, and editorials on the topic of brain-behavior associations and their reproducibility, related to Marek *et al.*, 2022. See the up-to-date list here: https://spisakt.github.io/BWAS_comment/

| Authors | Title | Where |
|---|---|---|
| Nature editorial | Cognitive neuroscience at the crossroads | Nature |
| Spisak et al. | Multivariate BWAS can be replicable with moderate sample sizes | Nature |
| [Nat. Neurosci. editorial ] | Revisiting doubt in neuroimaging research | Nat. Neurosci. |
| Monica D. Rosenberg and Emily S. Finn | How to establish robust brain–behavior relationships without thousands of individuals | Nat. Neurosci. |
| Bandettini P et al. | The challenge of BWAS: Unknown Unknowns in Feature Space and Variance | Med |
| Gratton C. et al. | Brain-behavior correlations: Two paths toward reliability | Neuron |
| Cecchetti L. and Handjaras G. | Reproducible brain-wide association studies do not necessarily require thousands of individuals | psyArXiv |
| Winkler A. et al. | We need better phenotypes | brainder.org |
| DeYoung C. et al. | Reproducible between-person brain-behavior associations do not always require thousands of individuals | psyArXiv |
| Gell M et al. | The Burden of Reliability: How Measurement Noise Limits Brain-Behaviour Predictions | bioRxiv |
| Tiego J. et al. | Precision behavioral phenotyping as a strategy for uncovering the biological correlates of psychopathology | OSF |
| Chakravarty MM. | Precision behavioral phenotyping as a strategy for uncovering the biological correlates of psychopathology | Nature Mental Health |
| White T. | Behavioral phenotypes, stochastic processes, entropy, evolution, and individual variability: Toward a unified field theory for neurodevelopment and psychopathology | OHBM Aperture Neuro |

| | | |
|---|---|---|
| Bandettini P. | Lost in transformation: fMRI power is diminished by unknown variability in methods and people | OHBM Aperture Neuro |
| Thirion B. | On the statistics of brain/behavior associations | OHBM Aperture Neuro |
| Tiego J., Fornito A. | Putting behaviour back into brain–behaviour correlation analyses | OHBM Aperture Neuro |
| Lucina QU. | Brain-behavior associations depend heavily on user-defined criteria | OHBM Aperture Neuro |
| Valk SL., Hettner MD. | Commentary on 'Reproducible brain-wide association studies require thousands of individuals' | OHBM Aperture Neuro |
| Kong XZ., et al. | Scanning reproducible brain-wide associations: sample size is all you need? | Psychoradiology |
| J. Goltermann, et al. | Cross-validation for the estimation of effect size generalizability in mass-univariate brain-wide association studies | BioRxiv |
| Kang K., et al. | Study design features that improve effect sizes in cross-sectional and longitudinal brain-wide association studies | BioRxiv |
| Makowski C., et al. | Reports of the death of brain-behavior associations have been greatly exaggerated | BioRxiv |
| J. Wu et al. | The challenges and prospects of brain-based prediction of behaviour | Nat. Human Behaviour |
